# T-toxin virulence genes: unconnected dots in a sea of repeats

**DOI:** 10.1101/2023.02.06.527415

**Authors:** Sajeet Haridas, Jennifer B. González, Robert Riley, Maxim Koriabine, Mi Yan, Vivian Ng, Adriana Rightmyer, Igor V. Grigoriev, Scott E. Baker, B. Gillian Turgeon

## Abstract

In 1970, the Southern Corn Leaf Blight epidemic ravaged US fields to great economic loss. The outbreak was caused by never-before-seen, super-virulent, Race T of the fungus *Cochliobolus heterostrophus*. The functional difference between Race T and O, the previously known, far less aggressive strain, is production of T-toxin, a host-selective polyketide. Super-virulence is associated with ∼1 Mb of Race T-specific DNA; only a fraction encodes T-toxin biosynthetic genes (*Tox1*). *Tox1* is genetically and physically complex, with unlinked loci (*Tox1A, Tox1B*) genetically inseparable from breakpoints of a Race O reciprocal translocation that generated hybrid Race T chromosomes. Previously, we identified ten genes for T-toxin biosynthesis. Unfortunately, high depth, short-read sequencing placed these genes on four small, unconnected scaffolds surrounded by repeated A+T rich sequence, concealing context. To sort out *Tox1* topology and pinpoint the hypothetical Race O translocation breakpoints corresponding to Race T-specific insertions, we undertook PacBio long-read sequencing which revealed *Tox1* gene arrangement and the breakpoints. Six *Tox1A* genes are arranged as three small islands in a Race T-specific sea (∼634 kb) of repeats. Four *Tox1B* genes are linked, on a large loop of Race T-specific DNA (∼210 kb). The race O breakpoints are short sequences of race O-specific DNA; corresponding positions in race T are large insertions of race T-specific, A+T rich DNA, often with similarity to transposable (predominantly Gypsy) elements. Nearby, are ‘Voyager Starship’ elements and DUF proteins. These elements may have facilitated *Tox1* integration into progenitor Race O and promoted large scale recombination resulting in race T.

**Importance:** In 1970 a corn disease epidemic ravaged fields in the US to great economic loss. The outbreak was caused by a never-before seen, super-virulent strain of the fungal pathogen *Cochliobolus heterostrophus*. This was a plant disease epidemic, however, the current COVID-19 pandemic of humans is a stark reminder that novel, highly virulent, pathogens evolve with devastating consequences, no matter what the host-animal, plant, or other organism. Long read DNA sequencing technology allowed in depth structural comparisons between the sole, previously known, much less aggressive, version of the pathogen and the super-virulent version and revealed, in meticulous detail, the structure of the unique virulence-causing DNA. These data are foundational for future analysis of mechanisms of DNA acquisition from a foreign source.

## Introduction

In 1970 an epidemic of Southern Corn Leaf Blight (SCLB) disease ravaged fields of corn on the eastern US seaboard to great economic loss. The outbreak was caused by a novel, super-virulent strain, Race T, of the fungal maize pathogen *Cochliobolus heterostrophus* (*Bipolaris maydis*). Although this was a plant disease epidemic, the current COVID-19 pandemic of humans reminds us that evolution of previously unseen pathogens, mechanisms of dissemination, and principles of disease are universal. Moreover, *C. heterostrophus* is particularly amenable to genetic and genomic functional analyses (1, 2) making these overarching questions more easily addressed than with some pathogens of animals for which host virulence assays are more burdensome.

The essential difference between *C. heterostrophus* Race T and Race O, the only previously known, but much less aggressive strain, is production by Race T of T-toxin, a polyketide metabolite not produced by Race O (3). Notably, all genes responsible for T-toxin biosynthesis are encoded in approximately one Megabase (Mb) of DNA sequence of Race T-specific DNA and super virulence is entirely attributable to this additional DNA (collectively known as *Tox1*) (4-7). Unlike most gene cohorts required for biosynthesis of natural products in filamentous ascomycete fungi (8), the *Tox1* genes are not tightly clustered (9). Rather, *Tox1* is both genetically and physically multifaceted. Decades of research have shown that *Tox1* genes are encoded at two unlinked loci (*Tox1A, Tox1B*) that are genetically inseparable from the breakpoints of a reciprocal translocation that formed a pair of hybrid Race T chromosomes (chromosomes 6;12, 12;6) when a pair of progenitor Race O chromosomes (chromosomes 6, 12) underwent a translocation (9, 10). The two Race T and two Race O chromosomes form a four-armed genetic linkage group with *Tox1* mapping to the intersection of this cruciform structure (4). At meiosis, progeny of a cross between Race T and Race O strains segregate 1:1 for ability to produce T-toxin, indicative of a single genetic locus, although we know two unlinked loci on two different chromosomes are involved.

Previously, ten *Tox1* genes required for biosynthesis of T-toxin were identified (9, 11). Genetic mapping, functional genetics, and in depth, short-read sequencing revealed that six of these genes are located at *Tox1A* and four at *Tox1B*, on four small, physically unconnected, genomic scaffolds. Three of the four scaffolds map to Race T chromosome 12;6 and the other to chromosome 6;12 (9). The *Tox1* genes are not found in Race O or in any of the more than 50 *Cochliobolus* species, however, as we reported recently (9), we have discovered orthologs of most *Tox1* genes as single, compact gene clusters in the genomes of three unrelated Eurotiomycetes and seven unrelated Dothideomycetes of various lifestyles, including plant pathogens, a mycoparasite, and two aquatic fungi, leading us to hypothesize that the tight gene cluster arrangement is ancestral. Importantly, in Condon et al (see Fig 1 in (9)), we could not determine *C. heterostrophus Tox1* gene order or the evolutionary progenitor. Thus, origin of the disjointed *Tox1* genes in *C. heterostrophus* remains unknown.

**Figure 1.**
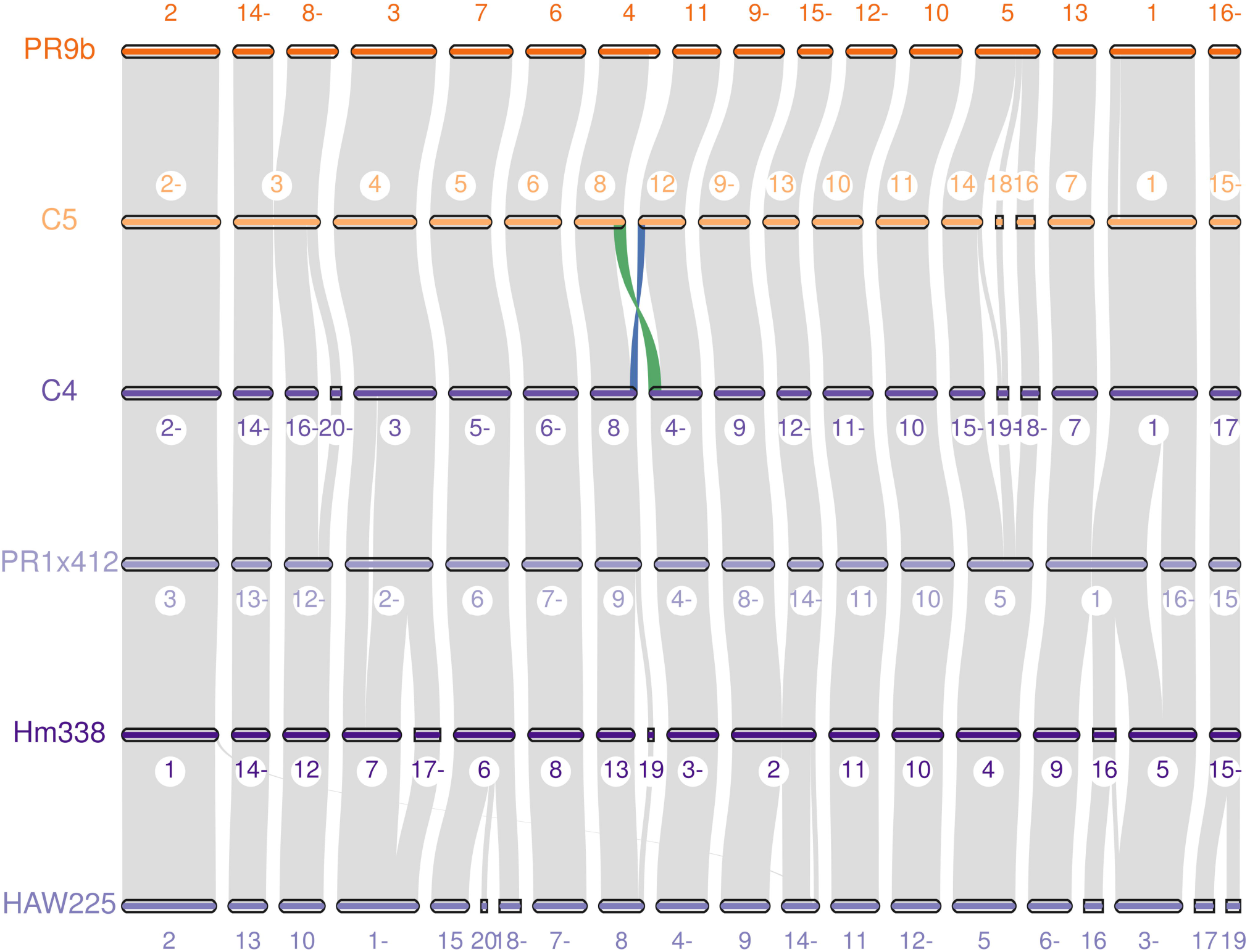
Macrosynteny visualization of six strains of *C. heterostrophus*. Race O strains are shown in orange while Race T strains are shown in purple. The numbers above the scaffolds in Race O and below in Race T, are scaffold numbers in the assembly. A minus sign after the scaffold numbers indicates opposite orientation to the sequence in the genome assembly fasta file. The blue and green ribbons show the translocation between chromosomes 6 and 12 in the Race T strains. Image made using jcvi mcscan (33).

To fully characterize the physical topology of *Tox1*-associated genes and scaffolds and identify the hypothetical translocation breakpoints in Race O, and the actual breakpoints in Race T chromosomes, which should correspond to points of insertion of Race T-specific DNA, we undertook long read sequencing of a collection of Race T and O strains (Table S1). Long read sequencing successfully spanned the translocation breakpoints in most strains, allowing identification of the breakpoints, the insertion points of Race T-specific DNA, and precise physical arrangement of *Tox1* genes. We found that, despite the fact that the six *Tox1A* genes are inseparable genetically, they are physically separated and arranged in tightly linked pairs at increasing distance from the genetic intersection of the four-armed linkage group. In total, *Tox1A* (known genes totaling 18.9 kb) consists of 634 kb of Race T-specific DNA in which three pairs of genes are islands in a sea of repetitive sequences. Four genes lie within 455 kb of Race T-specific sequence and two are found on 179 kb of Race T-specific sequence, separated from the former by 5.5 kb of sequence common to both Race T and Race O. The four genes at *Tox1B* (totaling ∼ 11 kb) are tightly linked genetically and physically, but also on a large stretch (210 kb) of Race T-specific DNA.

The theoretical breakpoint positions on the Race O chromosomes, which correspond to large insertions of Race T-specific DNA on the hybrid Race T chromosomes, are short stretches of Race O-specific DNA (2.3 kb/315 bp and 1.6 kb) on chromosomes 12 and 6, respectively. At these positions, Race T carries large inserts of Race T-specific DNA (179 kb, 455 kb on chromosome 12;6, 210 kb on chromosome 6;12) encoding *Tox1A* and *Tox1B* genes, respectively. These inserts are flanked on one side, by sequences homologous to Race O chromosome 6, and on the other by sequences homologous to Race O chromosome 12. Thus, the complex topological arrangement of all *Tox1* genes and the breakpoints of the reciprocal translocation are finally determined 50 years after the epidemic.

Additionally, the *Tox1* genes are embedded in blocks of repetitive A+T rich DNA, with many fragments that have homology to retrotransposons, particularly members of the Gypsy group. Nearby these repetitive DNA blocks, we found the sequence AGAATACAGGGC (GCCCTGTATTCT), which has been reported to be the target site (Voyager target site, vts) for the Voyager Starship element mediated by proteins DUF3435 and DUF3723 (12). We evaluate whether these may have facilitated integration of *Tox1* (source unknown) into a progenitor Race O strain.

## Results

### PacBio assemblies

While multiple efforts have been made to sequence *C. heterostrophus* Race O and Race T strains using both Sanger and Illumina short read technologies, the A+T rich and repetitive nature of the *Tox1A* and *Tox1B* loci severely limited efforts to assemble the region in its entirety. Although the majority of genic regions of the genome were captured, context was lost. Long read sequencing technologies significantly improved the fragmented nature of these assemblies, allowing complete construction of the *Tox1* regions. This is exemplified by the assemblies of both C4 (Race T) and C5 (Race O) near isogenic strain genome sequences (13-15) which improved from 207 to 70 and 68 to 53 scaffolds, respectively, and allowed more accurate gene coding and structural contextual comparisons (Fig. 1, Tables 1, S1-2). Assemblies generated from long read sequencing of field isolates of Race T and Race O strains have similar continuity. Most importantly, the new assemblies allow for deeper analysis of the multifaceted *Tox1* locus.

**Table 1.** Genome statistics for PacBio assemblies

| Assembly Statistic<sup>1</sup> | Race O |  |  | Race T |  |  |
| --- | --- | --- | --- | --- | --- | --- |
|  | PR9b | C5 | C4 | PR1x412 | Hm338 | HAW225 |
| Assembly Size (Mbp) | 38.41 | 36.48 | 37.79 | 41.32 | 45.25 | 44.2 |
| # Scaffolds | 113 | 53 | 70 | 67 | 133 | 111 |
| N50 & L50 (Mbp) | 7<br>(2.27) | 6<br>(2.12) | 7<br>(2.06) | 7<br>(2.23) | 9<br>(2.13) | 9<br>(1.92) |
| % Repeats | 14.97 | 11.46 | 13.23 | 18.31 | 21.05 | 23.63 |
| # Genes | 11,145 | 11,808 | 11,324 | 11,792 | 12,842 | 11,727 |

<sup>1</sup>https://mycocosm.jgi.doe.gov/Bipolaris_maydis

### Identification of *Tox1*-associated scaffolds

Because of the reciprocal translocation, flanking RFLP markers on the translocation-associated chromosomes differ between races T and O (Figs. 2, S1)(4). Race O chromosome 12 carries RFLP markers G395 and B88 which flank the theoretical breakpoint on this chromosome. Race O chromosome 6 carries G349 and B149 flanking the theoretical breakpoint. Race T hybrid chromosomes 12:6 and 6;12 carry G395/B149 and G349/B88, respectively. Using a BLAST-based approach with *Tox1* genes and flanking RFLPs as queries we identified the relevant pairs of markers on large single scaffolds in both Race O and Race T and thus captured the *Tox1* genes in Race T and reciprocal translocation breakpoints in both races (Figs. 2, S1, Table S3).

**Figure 2.**
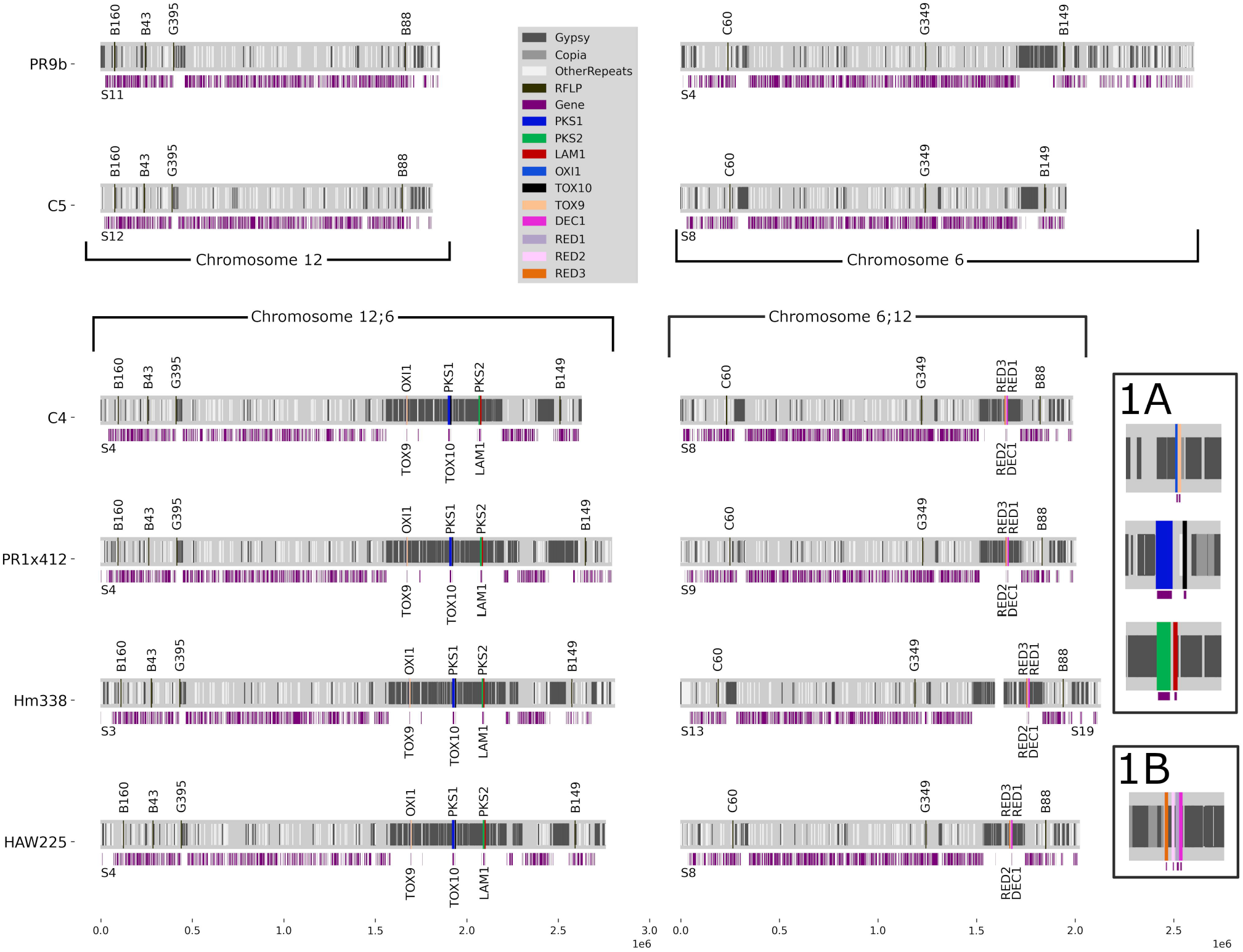
Diagram to scale of PacBio assembled Race O chromosomes 12 and 6 and Race T hybrid chromosomes 12;6 and 6;12 of all sequenced strains. Strains are listed on the left, color code for *Tox1* genes, RFLPs, and repeated elements on the right. Coordinates for the relevant scaffolds, *Tox1* genes and RFLPs listed in Table S3. Note Fig. 1 legend re a minus sign after a scaffold number. Automatically called genes are in purple beneath all chromosomes. Note absence of genes, except *Tox1* genes, in the repeat dense regions. Scale in Mb on the bottom.

For Race O near-isogenic strain C5, flanking RFLP marker pairs G395/B88 and G349/B149 were identified on single scaffolds S12 (1.811 Mb, chromosome 12) and S8 (1.952 Mb, chromosome 6), respectively, indicating complete sequencing through the hypothetical breakpoints on chromosomes 12 and 6 (Fig. 2, Table 2). For Race O field strain PR9b, flanking RFLP marker pair G395/B88 was identified on scaffold S11 (1.849 Mb, chromosome 12), while marker pair G349/B149 was identified on scaffold S4 (2.601 Mb, chromosome 6).

**Table 2.**
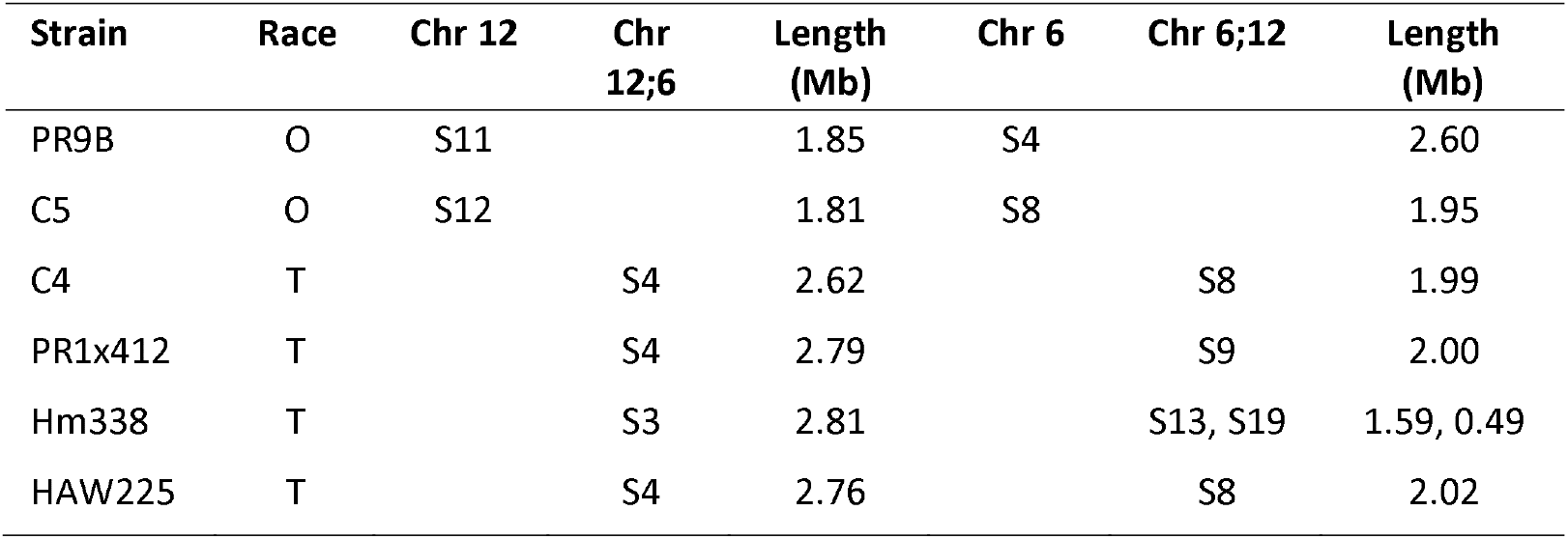
Mapping *Tox1* genes and RFLPs to scaffolds associated with Race O and T reciprocally translocated chromosomes

| Strain | Race | Chr 12 | Chr 12;6 | Length (Mb) | Chr 6 | Chr 6;12 | Length (Mb) |
| --- | --- | --- | --- | --- | --- | --- | --- |
| PR9B | O | S11 |  | 1.85 | S4 |  | 2.60 |
| C5 | O | S12 |  | 1.81 | S8 |  | 1.95 |
| C4 | T |  | S4 | 2.62 |  | S8 | 1.99 |
| PR1x412 | T |  | S4 | 2.79 |  | S9 | 2.00 |
| Hm338 | T |  | S3 | 2.81 |  | S13, S19 | 1.59, 0.49 |
| HAW225 | T |  | S4 | 2.76 |  | S8 | 2.02 |

For Race T near-isogenic strain C4, the *Tox1A* genes, previously identified on three small unconnected scaffolds in the short read assemblies, were all on single scaffold S4 (2.625 Mb, chromosome 12;6) in the PacBio assembly, flanked by RFLP markers G395 and B149. In Race T field strains PR1×412, Hm338, and HAW225, *Tox1A* and the same flanking G395/B149 markers were also located on single scaffolds (PR1×412: S4, 2.791 Mb; Hm338: S3, 2.807 Mb; HAW225: S4, 2.757 Mb, chromosome 12;6) (Fig. 2, Table 2).

For Race T *Tox1B* genes, we resolved chromosome 6;12 for strains C4, PR1×412, and HAW225, where RFLP G349, *Tox1B*, and RFLP B88 map to single scaffolds (C4: S8, 1.987 Mb; PR1×412: S9 (2.0 Mb, HAW225: S8, 2.021 Mb, chromosome 6;12) (Fig. 2, Table 2). For strain Hm338, *Tox1B* genes and RFLP marker B88 were located on single scaffold S19 (0.488 Mb), but RFLP G349 was on separate scaffold S13 (1.590 Mb).

Thus, the entire *Tox1A* and *Tox1B* loci and their flanking RFLP markers have been captured in all Race T (with one exception at *Tox1B*) and both Race O strains, allowing an in-depth examination of differences between the races at the *Tox1* locus.

### Mapping the *Tox1* genes-tightly linked genetic distances, long physical distances

In the short read assembly of strain C4 (v1), the 10 known *Tox1* genes mapped to four very small, unconnected, scaffolds. Three scaffolds carry *Tox1A* genes which mapped in pairs on separate scaffolds: *PKS1*/*TOX10* (S74, ∼17 kb), *PKS2*/*LAM1* (S83, ∼11 kb) and *OXI1*/*TOX9* (S102, ∼3.5 kb). The fourth carries *Tox1B* genes in a tight cluster, also on a small scaffold (S82, ∼13.5 kb). The PacBio assembly of four Race T strains, including inbred lab strain C4, indicates that all six genes at *Tox1A* are captured on single scaffolds, arranged in the order *LAM1, PKS2, TOX10, PKS1, TOX9, OXI1*, in tightly linked pairs of increasing distance, respectively, from the genetic intersection of the four-armed linkage group (Figs. 2, 3, S1). In total, *Tox1A* consists of 634 kb of Race T-specific DNA in which the three pairs of genes (totaling 18.9 kb) are islands in a sea of repetitive sequences. *PKS2/LAM1, PKS1/TOX10* are on 455 kb of Race T-specific sequence and *OXI1/TOX9* is on 179 kb of Race T-specific sequence, separated from the former by 5.5 kb of sequence common to both T and O races. The *PKS1/TOX10* pair is 225 kb from *OXI1/TOX9*. The *PKS2/LAM1* pair is 153 kb away from the *PKS1/TOX10* pair (Fig. 3). Note, furthermore, although genetic mapping indicated that the six genes at *Tox1A* are genetically inseparable from the reciprocal translocation breakpoints, they are, in fact, physically separated from each other by hundreds of kilobases and not in a cluster like most secondary metabolite-associated genes. Interestingly, in all Race T strains, the two large loops of Race T-specific *Tox1A* DNA are separated by 5.5 kb of DNA common to both Race T and O.

**Figure 3.**
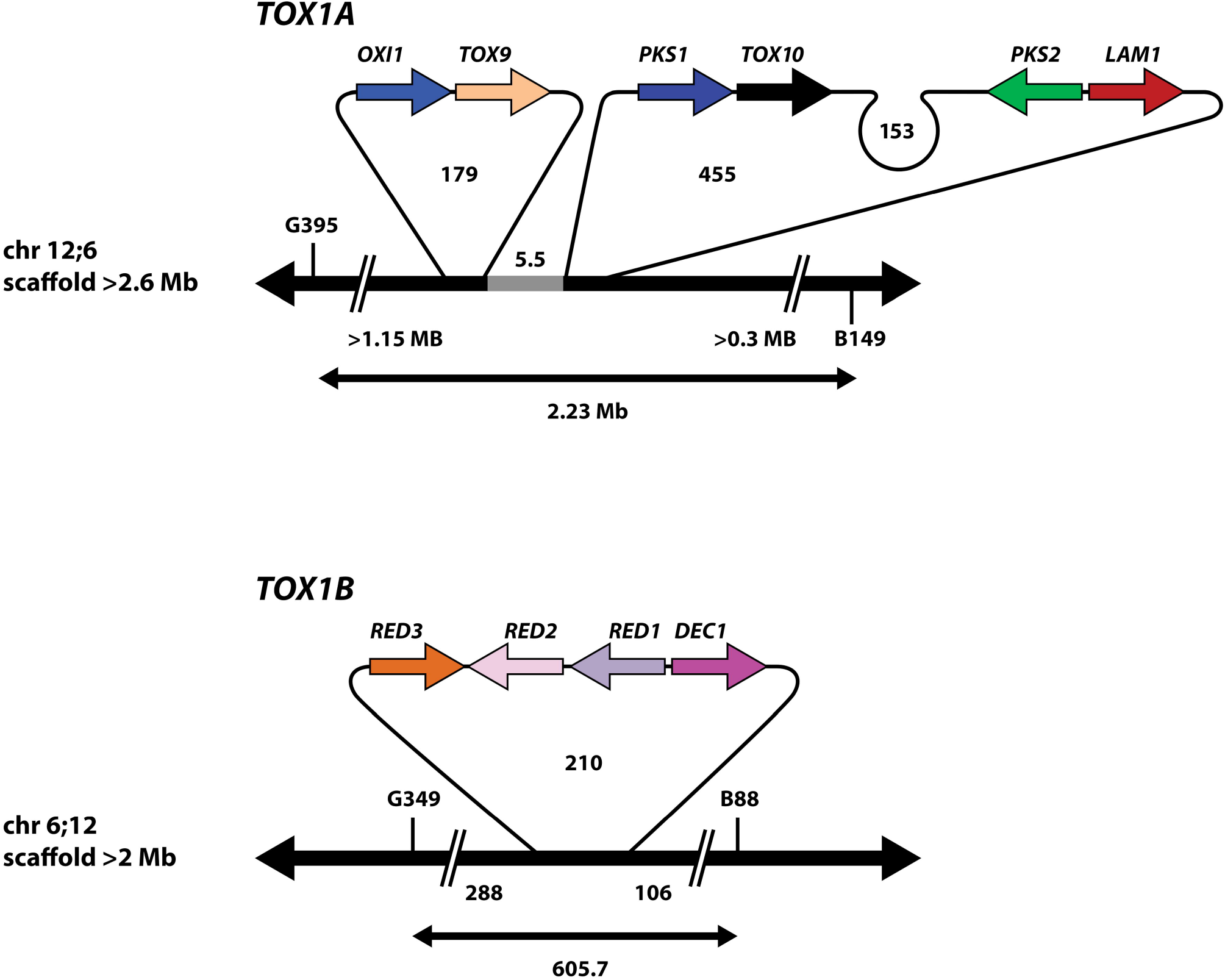
Structural topography of the *Tox1* region. The exact order and spatial arrangement of known *Tox1* genes is shown. There are six genes at *Tox1A*, which are found as three tightly linked pairs of genes, separated by 100s of kb of A+T rich DNA. *OXI1/TOX9* is on a 179 kb loop separated from a 455 kb loop carrying *PKS1/TOX10* and *PKS2/LAM1*. 5.5 kb of DNA (light gray) in common to both Race T and Race O separates the *Tox1A* loops. *OXI1* is furthest from the breakpoint while *LAM1* is closest on chromosome 12:6. There are four genes at *Tox1B* clustered tightly together on chromosome 6;12 on a 210 kb loop of A+T rich DNA. All numbers are in kb except where indicated (not to scale). Double-headed arrows below each locus is distance between RFLP marker pairs G395/B149 (*Tox1A*) or G349/B88 (*Tox1B*), respectively. >1.15Mb, >0.3 Mb, 288, and 106 are distances from the nearest RFLP marker to the beginning of the nearest loop.

As described above, the *Tox1A* genes are found in pairs separated from each other by vast stretches of Race T-specific DNA that is A+T rich, highly repeated, and devoid of genes apart from the *Tox1* genes (Figs. 2, 3). Exceptions include two genes in the 5.5 kb DNA in common to both races, between the Race T *Tox1A* specific loops, which encode a predicted short chain dehydrogenase and an alpha/beta hydrolase, corresponding to strain C5 (v3) JGI protein IDs 1196534 and 1226739, respectively. These two genes are highly conserved in most Dothideomycetes and therefore not unique to Race T, as are all known *Tox1* genes. The two genes occur adjacent to each other in all *Cochliobolus* species for which we have sequence, suggesting conservation across the species. Furthermore, the genes flanking these two genes (C5 v3 JGI protein IDs 2337044 and 1142280) in race O, are found on two different chromosomes (S8, chromosome 12;6 and S4 chromosome 6;12, respectively) in race T as predicted, reflective of their change in position due to the translocation. This places the theoretical reciprocal translocation breakpoint in race O on chromosome 12 between genes encoding proteins 2337044 and 1226739 (or ∼ within 1118 bp), which overlaps with the position of the 315 bp race O-specific DNA (Fig. S2).

The four *Tox1B* genes are tightly clustered, as noted earlier (9, 15, 16). Remarkably, no additional genes were predicted in the 210 kb Race T-specific loop (Figs. 2, 3).

Thus, the topographical organization and exact genomic position of the previously unconnected *Tox1A* genes were clarified by long read sequencing for the first time. This approach also extended the flanking A+T replete sequence surrounding *Tox1A* and *Tox1B* encoding genes and revealed these are sizeable expanses compared to the small tracts of Tox1 coding sequence. The A+T rich blocks on Tox1-associated scaffolds (Fig. 2) are replete with gypsy element fragments (most are ∼75% A+T), a few of which are full open reading frames with high identity to multiple open reading frames in both Race T and Race O genomes suggesting these are native to *C. heterostrophus*. We speculate that their presence in Tox1-associated DNA is due to movement in Race T after the introduction of Tox1 DNA (still unknown donor) (9).

### *Tox1*-associated scaffolds from geographically dispersed Race T and Race O strains are homologs

Diagnostic RFLPs (G395/B149, G349/B88) and *Tox1* genes map to chromosomes 12;6 and 6;12 in all Race T strains (Fig. 2, Table 2). Similarly, the diagnostic RFLP pairs (G395/B88, G349/B149) flanking the theoretical breakpoints in Race O map to chromosomes 12 or 6. This reflects conservation of the reciprocal translocation, the Race O theoretical breakpoints, and insertion points of *Tox1* DNA in strains of wide geographical distribution. Furthermore, for Race T chromosome 12:6, the *Tox1A* genes are conserved in order, gene orientation, and position (Fig. 3). Distance differences between flanking RFLP markers G395 and B149, are likely due to variations in the number of repeat sequences encoding transposable elements associated with the Race T-specific loops (Fig. 2).

Our findings that *Tox1*-associated scaffolds from Race T strains are syntenic and that the *Tox1* genes are identically arranged, support the hypothesis that Race T arose once from a single genetic event, at an unknown time, from an, as yet unknown source, in which *Tox1* DNA integrated into a chromosome of a Race O strain. This strengthens earlier genetic and molecular karyotype data (17) which indicated all T strains carried the reciprocally translocated chromosomes.

### The breakpoints of the reciprocal translocation

Race O inbred strain C5 and Race T inbred strain C4 and Race O inbred strain C5 and Race T field isolate PR1×412 were chosen as representative isolates for further analysis (one Race O strain compared to one Race T inbred and one field strain). Relevant *Tox1* genes and RFLP markers are on single continuous scaffolds in all (Fig. 2). Circos plots of strain C5 chromosomes 12 and 6 and strain PR1×412 or C4 chromosomes 12;6 and 6;12 support the reciprocal translocation hypothesis in showing that Race T chromosomes 12;6 and 6;12 each have homology to both Race O chromosomes 6 and 12, hence providing additional evidence at the nucleotide level for the genetically and physically documented reciprocal translocation (Fig. 4). For chromosome 12;6, the largest portion of identity is with Race O chromosome 12, while for chromosome 6;12, the largest portion of identity is with Race O chromosome 6. Both physical findings support the genetic convention of naming hybrid chromosomes according to the largest portion contributing to the hybrid, first.

**Figure 4.**
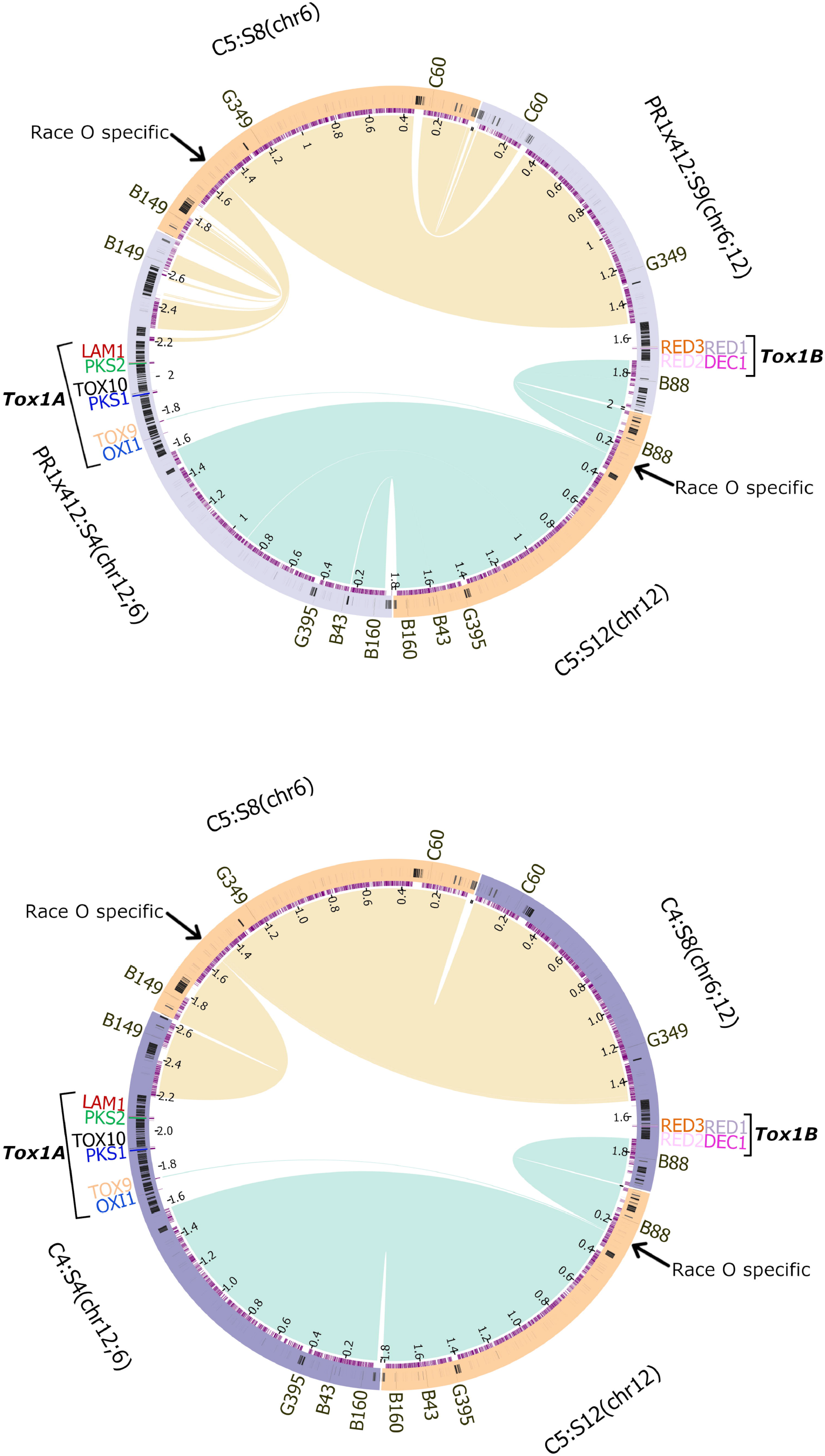
Circos diagrams of C5 (Race O) chromosomes 6 and 12 and the reciprocally translocated PR1×412 or C4 (Race T) chromosomes 12;6 and 6;12 counterparts. Each Race O chromosome has a match to both Race T hybrid chromosomes. On Race O chromosome 6, the arrow points to 1.6 kb of Race O-specific DNA whose counterpart on chromosome 6;12 is the 210 kb *Tox1B* Race T-specific insert. On Race O chromosome 12, the arrow points to 315 and 2.3 kb of Race O-specific DNA, separated by 5.5 kb of DNA common to both races. The 315 Race O-specific DNA counterpart position in Race T is the 455 kb insert of *Tox1A* Race T-specific DNA carrying *PKS2/LAM1* and *PKS1/TOX10*, while the 2.3 kb Race O counterpart is the 179 kb *Tox1A* insert carrying *OXI1/TOX9*, on chromosome 12;6.

The Circos plots also illustrate that both theoretical breakpoints in Race O correspond to large inserts of Race T-specific DNA on chromosomes 12;6 and 6;12 (Fig. 4, Table S4). Race O strain C5 chromosome 12 (scaffold 12) at the arrow in Fig. 4, actually corresponds to two Race O specific stretches of DNA (315 bp and 2.3 kb, respectively, Fig. 5), separated by 5.5 kb of DNA in common between the two races (Fig. S2). At these positions, *Tox1A* carries Race T-specific inserts of 179 kb encoding the *OXI1/TOX9* gene island and 455 kb encoding the *PKS1/TOX10* and *PKS2/LAM1* islands, separated by 5.5 kb of DNA in common between the two races. We speculate that this 5.5kb island of synteny between the O and T strains, which splits *Tox1A* is indicative of an additional recombination event, post initial insertion event. Chromosome 6 (scaffold 8) at the arrow in Fig. 4, corresponds to a Race T specific insert of 210 kb, carrying the tightly linked Tox1B genes (*DEC1/RED1/RED2/RED3*) (Figs. 4, 5, Table S4).

**Figure 5.**
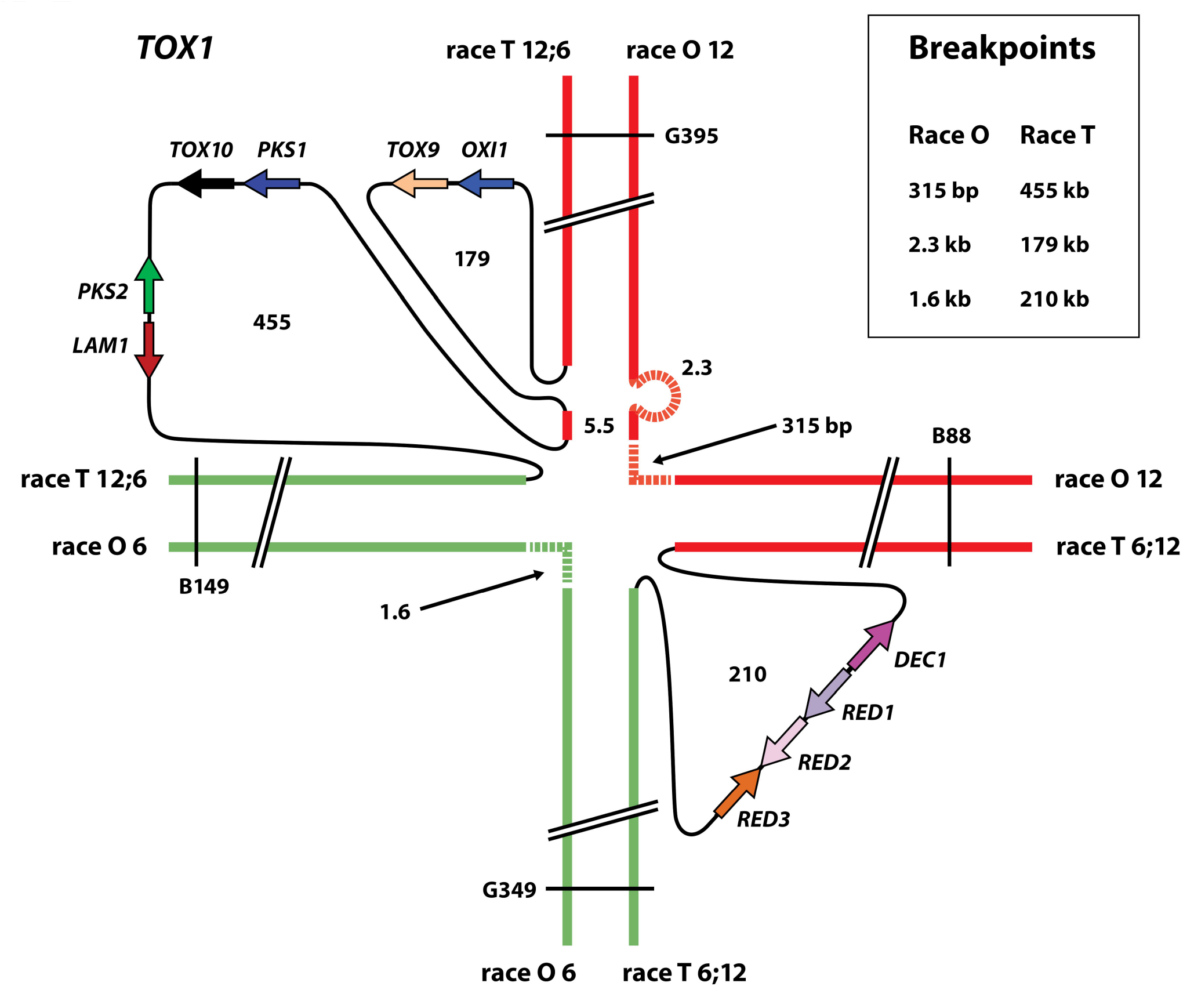
Model of the complex genetic structure of the *Tox1* locus and linkage map of chromosomes involved in the reciprocal translocation. Race O chromosome 12 in red, chromosome 6 in green. Race T hybrid chromosomes in red and green. Black loops indicate inserts of Race T-specific DNA corresponding to positions of Race O-specific DNA (dotted lines) as summarized in box insert. Figure is not to scale. All numbers in kb, except where indicated. Key RFLPs are shown (Fig. S1).

Remarkably, the theoretical Race O reciprocal translocation breakpoints are not sequences that are split and rearranged in Race T, but rather short stretches of Race O-specific DNA. These consist of 315 bp and 2.3 kb of sequence on chromosome 12 and 1.6 kb of sequence on chromosome 6 (Figs. 4, 5 Table S4). As noted above, large inserts of Race T-specific DNA are found at the corresponding positions on Race T chromosomes 12;6 and 6;12 flanked by DNA from either Race O chromosome 6 or 12. The repeat islands around *Tox1A* locus are significantly larger than those around *Tox1B*.

## Discussion

Facilitated by long-read sequencing, megabase scale, nucleotide-level resolution has revealed the true organization of *Tox1*, the locus responsible for T-toxin production and the SCLB epidemic of 1970. Four Race T strains, collected shortly after the epidemic, but representing wide geographical distribution, were sequenced. We found that all *Tox1*-associated scaffolds/chromosomes were homologs, and the 10 known *Tox1* gene positions, order, and direction were identical in all. We also determined, in every case, that the known unlinked *Tox1A* and *Tox1B* genes are small islands in a sea of Race T-specific repeats and that many sequences with similarity to A+T rich Gypsy transposons are found in this sea. These findings are strong verification not only for the reciprocal translocation but also the hypothesis that a one-time complex genetic event gave rise to Race T from a progenitor Race O strain.

As we reported in Condon et al (9), orthologs of most *Tox1* genes have been found as single, compact gene clusters in the genomes of three unrelated Eurotiomycetes and seven unrelated Dothideomycete species, which led us to hypothesize that the tight gene cluster arrangement is ancestral and that these species likely produce a metabolite that differs from T-toxin. Note that in Condon et al (9), (see Fig. 1), we could not even determine *C. heterostrophus* Tox1 gene order because the genes were distributed on four very small scaffolds, which we could not connect by any means. Comparative analyses showed that only *C. heterostrophus* and *Didymella zeae-maydis* carry *RED1*, an NAD dependent oxidoreductase. Notably, *D. zeae-maydis* produces PM-toxin, a linear polyketide toxin, similar to T-toxin, and was also discovered around the time of the SCLB epidemic as it affects the same host and causes similar devastation of T-cms corn (18-20). Phylogenetic analyses and similarity assays confirmed that neither of these taxa passed *Tox1* to the other (9). In addition, most of the tight clusters in the other fungi encode an ABC transporter and an oxidoreductase in the middle of their clusters, neither of which are found anywhere in either the *C. heterostrophus* Race T or O genomes (Condon et al. (9), see Fig. 1). These genetic differences are likely key to functional differences, as not one of these *Tox1*-like carrying species produces a T-toxin like symptom using the T-toxin-specific microbial assay (9). The evolutionary progenitor of the disjointed *Tox1* genes in *C. heterostrophus* Race T remains unknown and is not our focus here. Rather, here, with long read sequencing, we have determined that the *C. heterostrophus Tox1* genes are in a different order from all other species known to carry *Tox1*-like genes, confirmed they are not clustered and revealed their context in large blocks of repeated A+T DNA. It must be acknowledged that all Race T isolates in repositories, including ours, represent strains collected worldwide within a few years of the SCLB epidemic. In our case, these strains have been stored on silica gel or as glycerol stocks at -80C and revitalized for each experiment, thus they are ‘frozen in time’. Yet, as noted below, the ability of Race T strains to produce T-toxin is stable in culture and long-term storage suggesting it is stable in the population, as reported earlier (17) but that Race T strains are not as fit as race O strains, unless there’s strong selection (21-23). Indeed, when T-cms corn, the specific host for supervirulent Race T and *D. zeae-maydis* was no longer planted, Race *T/D. zeae-maydis* disappeared from the field.

### Reciprocal translocation breakpoints

The Race O theoretical reciprocal translocation breakpoints were found not to be single nucleotide transition points, but rather stretches of Race O-specific DNA (315 bp and 2.3 kb on Race O chromosome 12, and 1.6 kb on chromosome 6). These breakpoints were evident in both Race O strains (field and inbred) that were sequenced by PacBio. Interestingly, the Race O-specific DNA and corresponding Race T inserts on chromosome 12 are interrupted by 5.5 kb of DNA common to both races. We speculate that one or more additional recombination events occurred in Race T after the initial *Tox1* DNA insertion and reciprocal translocation events and that this resulted in separation of the two *Tox1A* loops.

### Repeated regions

The cultivation of monocultures of Race T-susceptible T-cms maize and complex DNA recombination events including a reciprocal translocation are considered the primary causes of the SCLB epidemic that occurred in the 1970s (6). Were the vast blocks of associated repetitive DNA part of the original transfer? As reported earlier (24), ∼ 50% of the TE sequences identified in the Race O genome belong to the Gypsy superfamily of retrotransposons and it is known that retrotransposons are the largest constituent of the repetitive fraction of the genome in phytopathogenic fungi (25). At least some of these regions correspond to centromeres which, although A+T rich and often associated with transposable elements, vary widely in structure and size in fungi (26). Indeed, PacBio sequencing has confirmed that the sea of repetitive DNA surrounding the *Tox1* genes is Gypsy element replete. Some of these are full ORFs with high quality matches to multiple open reading frames in both Race T and Race O genomes which suggests they were already in the progenitor Race O genome before Race T appeared. It is possible they are/were active and perhaps were involved in proliferation of DNA within the Race T-specific *Tox1* blocks after insertion of *Tox1*.

Although we’ve speculated that the original insert of *Tox1* DNA was into a centromere, as these have characteristics similar to *Tox1* regions (repeats, A+T rich, transposable elements), we currently have no conclusive evidence for this, as we did not perform experiments to specifically locate centromeres in our race T and O strains. Chromosomal rearrangements, associated with transposable elements have been demonstrated recently for *Verticillium* (27), and *Aspergillus fumigatus* (28), as well as plant genomes (29).

### Are Starship elements involved?

We explored the possibility that *Tox1* may be a giant Starship element (12) and looked for diagnostic features of these elements around *Tox1*. Flanking the B149 RFLP marker (Figs. 2, S1) in both the O and T strains are genes encoding proteins with conserved pfam domains DUF3435 and DUF3723 thought to mediate integration into the Voyager Starship element target site (vts). Approximately 300 kb upstream of B149 (270kb - 380kb, depending on the assembly), we identified the vts sequence (AGAATACAGGGC) in Race O and also just outside the repeat block harboring the *Tox1A* locus in Race T (Fig. S3). In both races this is ∼240kb from DUF3435, whereas the average size of the region between vts and the DUF genes reported, originally, in (12) is ∼100kb. Note, the 12 bp vts sequence is rare in both race T and O genomes, seen only once on the scaffolds harboring B149.

We did not find an association with genes encoding 5S rDNA, into which Starship elements are reported to insert, or with other genes commonly localized with Starship elements (FRE, PLP, NLR), nor did we identify associated inverted terminal repeats (12). Therefore, although we found the vts, and two DUF proteins, we did not find other features commonly associated with Starship. We speculate, given the bizarrely complex organization of *Tox1*, that at least two massive recombination events occurred and that these events may have erased diagnostic signatures of Starships or other known mobile elements. We speculate that an investigation of the evolutionary history of Race O-specific gaps at the theoretical breakpoints will be revealing.

### Evolutionary mechanism

As reported in Condon et al (9) the genetic source of *C. heterostrophus Tox1*, responsible for biosynthesis of T-toxin and the ravaging SCLB epidemic, remains unclear. In all cases where they are found, most *Tox1* or *Tox1*-like genes have introns, thus, are likely of eukaryotic origin plus orthologs group together phylogenetically which suggests a common evolutionary origin. *Tox1* clusters in non-T-toxin producing species are in compact linear array, which is likely the ancestral arrangement. Here, we have used PacBio sequencing to advance understanding of *C. heterostrophus Tox1* gene topological organization and overall fine genetic structure >50 years after the epidemic. Evolutionary underpinnings defy understanding at this point. That said, we would argue that evolution of *Tox1* included an initial insertion of foreign, likely eukaryotic DNA, a reciprocal translocation, and one or more recombination events promoted by mobile native gypsy elements that caused amplification of the A+T rich sea associated with *Tox1* DNA.

The story of *C. heterostrophus*, T-cytoplasm corn and T-toxin is a timely reminder that natural selection drives both the rapid rise and decline of disease. One remarkable aspect of *C. heterostrophus* Tox1 is its high level of expression (30) and stability. We hypothesize that selective pressure from the widespread monoculture of T-cms maize either led to the acquisition of the biosynthetic cluster around the time of the epidemic or, alternatively, Race T might have been ‘lurking’ in the field and the T-cms maize monoculture made it an ideal ‘petri dish’ for selection for Race T. Strong selection provided ability to persist in the field amidst a background population of Race O; the latter continues even in progeny of laboratory crosses with Race O. In either case, these properties made *C. heterostrophus* a “chassis” for synthetic biology driven by natural selection in the environment.

## Methods

### Strains

*C. heterostrophus* strains sequenced by PacBio (Table S1) included highly backcrossed lab strains C5 (ATCC 48332, Race O, *MAT1-1, Tox1*-) and C4 (ATCC48331, Race T, *MAT1-2, Tox1+*), expected to differ significantly only at *Tox1* and *MAT1* (mating type) after 6 backcrosses to a recurrent parent. The other strains were field isolates of wide geographic origin, collected around the time of the 1970 SCLB epidemic and included one Race O field isolate PR9b (Poza Rica, Mexico, *MAT1-1*) and three Race T field isolates as follows: Hm338 (New York, *MAT1-2, Tox1+*, ATCC 48317), PR1×412, (a progeny of a cross between PR1C from Poza Rica, Mexico and strain 412, unknown geographical origin, Race T, *MAT1-1, Tox1+*) and HAW225 (Hawaii, *MAT1-1, Tox1+*). Aliquots of strains from -80°C glycerol stocks were plated on complete medium with xylose (CMX) for one week (16/8hr light, 23C) as previously described (31).

### High molecular weight DNA preparation

1000 mls of liquid CM medium were inoculated with 10^6^ -10^7^ spores collected from CMX plates and shaken at 150-200 rpm at room temperature for two days. Mycelium was harvested on Whatman #4 filter paper using a Buchner funnel connected to a vacuum. The mycelium pad was peeled off the filter, placed in a 50 ml Falcon tube, and stored at -20°C or directly lyophilized for 25 hrs (cap loose) or until mycelium was dry and brittle. Lyophilized mycelia were pulverized with a pre-chilled pestle in a pre-chilled mortar to a fine powder. Mycelial powder (∼0.2-0.5g) was suspended in 20ml isolation buffer (150 mM EDTA pH 8.0, 50 mM Tris, pH 8.0, 1 % (w/v) sarkosyl (n-lauroyl sarcosine), 300 mg/L Protease XI (Proteinase K) in disposable 30ml polypropylene tubes (Sarstedt) and vortexed vigorously for 30-60 sec. Cell debris was pelleted by centrifugation in an SS-34 rotor (5 min, 5000 rpm, 4°C) and supernatant transferred to a clean tube. 20ml isolation buffer was added and centrifugation repeated. The recovered supernatant was gently mixed with an equal volume of Tris-saturated phenol and tubes centrifuged (10 min, 5000 rpm, 4°C) to separate phases. The upper (aqueous) phase was transferred to a clean tube and mixed with an equal volume of 25:24:1 phenol:chloroform:isoamyl alcohol. Centrifugation and recovery of the top aqueous layer was repeated twice. Finally the aqueous layers were mixed with an equal volume of 24:1 chloroform:isoamyl alcohol and centrifuged as before. DNA was extracted by ethanol precipitation (1/10 vol. of 3M NaOAc plus, 2 vol. of cold absolute ethanol) for 10 min at -20°C and centrifuged as above. The pellet was transferred to a 1.5 ml tube, washed with 70% ethanol twice, centrifuged (5 min, 5K rpm, 4°C) and air dried. The DNA was dissolved in 2x 200ul TE (65°C 10-20 min to dissolve, check frequently, not longer than 30 min), 2 ul RNase A (10ug/ul, Sigma R-6513) was added to each tube and incubated at 37°C for 1 hr. A l.5ul aliquot was run on a 0.7% agarose gel to check if RNA was still present.

### Sequencing and annotation methods

For all *C. heterostrophus* genomes (except C5), 10 ug of genomic DNA was sheared to >10kb using Covaris g-Tubes. The sheared DNA was treated with exonuclease to remove single-stranded ends and DNA damage repair mix followed by end repair and ligation of blunt adapters using SMRTbell Template Prep Kit 1.0 (Pacific Biosciences). The library was purified with AMPure PB beads and size selected with BluePippin (Sage Science) at >10 kb cutoff size. PacBio Sequencing primer was then annealed to the SMRTbell template library and sequencing polymerase was bound to them using Sequel Binding kit 3.0. The prepared SMRTbell template libraries were then sequenced on a Pacific Biosystems Sequel sequencer using v3 sequencing primer, 1M v3 SMRT cells, and Version 3.0 sequencing chemistry with 1×600 sequencing movie run times. Filtered subread data was processed to remove artifacts and assembled with Falcon version pb-assembly=0.0.2|falcon-kit=1.2.3|pypeflow=2.1.0 (https://github.com/PacificBiosciences/FALCON) to generate an initial assembly. Mitochondrial scaffolds were identified in the initial assembly and reads mapping to this were filtered out. A secondary Falcon assembly was generated using the mitochondria-filtered preads, improved with finisherSC version 2.1 (32), and polished with Arrow version SMRTLINK v7.0.1.66975 (https://github.com/PacificBiosciences/GenomicConsensus).

10 ug of C5 genomic DNA was sheared to 30 kb - 50 kb using the Megaruptor 3 (Diagenode). The sheared DNA was treated with exonuclease to remove single-stranded ends, DNA damage repair enzyme mix, end-repair/A-tailing mix and ligated with overhang adapters using SMRTbell Express Template Prep Kit 2.0 (PacBio) and purified with AMPure PB Beads (PacBio). Individual libraries were size-selected using the 0.75% agarose gel cassettes with Marker S1 and High Pass protocol on the BluePippin (Sage Science). PacBio Sequencing primer was then annealed to the SMRTbell template library and sequencing polymerase was bound to them using Sequel II Binding kit 2.0. The prepared SMRTbell template libraries were then sequenced on a Pacific Biosystems Sequel II sequencer using tbd-sample dependent sequencing primer, 8M v1 SMRT cells, and Version 2.0 sequencing chemistry with 1×900 and 1×1800 sequencing movie run times. Filtered subread data was processed to remove artifacts, subsampled to 10% using BBtools (BBMap – Bushnell B. – sourceforge.net/projects/bbmap/) version 38.87 [reformat.sh samplerate=0.1] and assembled with Flye version 2.8.1-b1676 (https://github.com/fenderglass/Flye) to generate an assembly, which was polished with gcpp-- algorithm arrow version SMRTLINK v8.0.0.80529 (https://www.pacb.com/support/software-downloads).

For all assemblies, contigs less than 1000 bp were excluded. Further genome improvement was done using reads recruited to scaffold ends using bbtools (bbduk.sh), reassembled with Flye and polished with finisherSC. Some scaffolds were joined using reassembly of locally mapped reads.

Mitochondria was assembled separately from the Falcon pre-assembled reads (preads) with an in-house tool using the filtered preads and polished with Arrow version SMRTLink v7.0.1.66975. The C5 mitochondrial DNA was assembled separately with CS reads using an in-house tool and polished with gcpp --algorithm arrow version SMRTLink v8.0.0.80529.

Each genome was annotated using the JGI Annotation Pipeline (Grigoriev et al., 2014)

### Mapping of *Tox1* genes and RFLP markers

Nucleotide sequences for *Tox1* genes and selected RFLP markers associated with the *Tox1* locus at the intersection of the four-armed linkage group (B43, B88, B149, B160, C60, G349, G395, (4) shared between Race O and Race T chromosomes were used to identify *Tox1*-associated scaffolds in the PacBio assemblies using blast. (Fig. S1, Table S3).

## Supporting information

Sippl Fig. 2

## Data availability

Genome assemblies and annotations for the organisms used in this study are available via the JGI fungal genome portal MycoCosm (Grigoriev eta al., 2014; http://mycocosm.jgi.doe.gov) and have been deposited to GenBank under accessions XXXXXXXXX

### Acknowledgements

BGT would like to thank Olen Yoder for introducing her to the remarkable effects of T-toxin and all current and former lab members for their relentless enthusiasm for genetic and genomic discoveries. This work (proposal: 10.46936/10.25585/60001208) conducted by the U.S. Department of Energy Joint Genome Institute (https://ror.org/04xm1d337), a DOE Office of Science User Facility, is supported by the Office of Science of the U.S. Department of Energy operated under Contract No. DE-AC02-05CH11231. A portion of this work was part of the DOE Joint BioEnergy Institute (http://www.jbei.org) supported by the U. S. Department of Energy, Office of Science, Office of Biological and Environmental Research, through contract DE-AC02-05CH11231 between Lawrence Berkeley National Laboratory and the U. S. Department of Energy.

## Supplementary Tables

S1: Genotypes and sources of strains used in this study.

S2: Annotation statistics of the genomes used in this study.

S3: Coordinates of the RFLP markers and *Tox1* genes.

S4: Synteny data used in Fig 4.

## Figure legends

**Figure S1.** Genetic map of the four-armed linkage group associated with the *Tox1* locus.

Selected RFLP markers used in this study and their locations relative to *Tox1* (intersection) are shown (modified from (4, 34)). Race T chromosomes = thin solid line, Race O chromosomes = dashed lines. The RFLP markers = thin lines connecting chromosome lines. Thick broken line represents statistically linked regions of chromosomes. T6-2 and T12-1 are telomeres. (Note that the numbers 12 and 6 on the left of the semicolon indicate which Race O chromosomal counterpart is the largest fragment of the hybrid chromosome).

**Figure S2.** Screenshot of the breakpoint positions in Race O chromosome 12 and the corresponding positions in race T (https://mycocosm.jgi.doe.gov/cgi-bin/browserLoad/?db=CocheC5_4m&position=scaffold_12:252553-282945). The common 5.5 kb region is indicated (Figs. 3, 5) flanked by sequences that occur only in Race O (red arrows). Strain C5 v3 protein numbers corresponding to predicted genes are indicated.

**Figure S3.** Diagram of PacBio assembled chromosomes.

Race O chromosomes 12 and 6 and Race T chromosomes 12;6 and 6;12 of all sequenced strains indicating locations of candidate Starship elements are shown. See Fig. 1. Vts, DUF3435 and DUF3723 are shown relative to RFLP marker B149 in both races and *Tox1* in Race T.

