## Supplementary figures and images for "T-toxin virulence genes: unconnected dots in a sea of repeats"

### Sippl Fig. 2

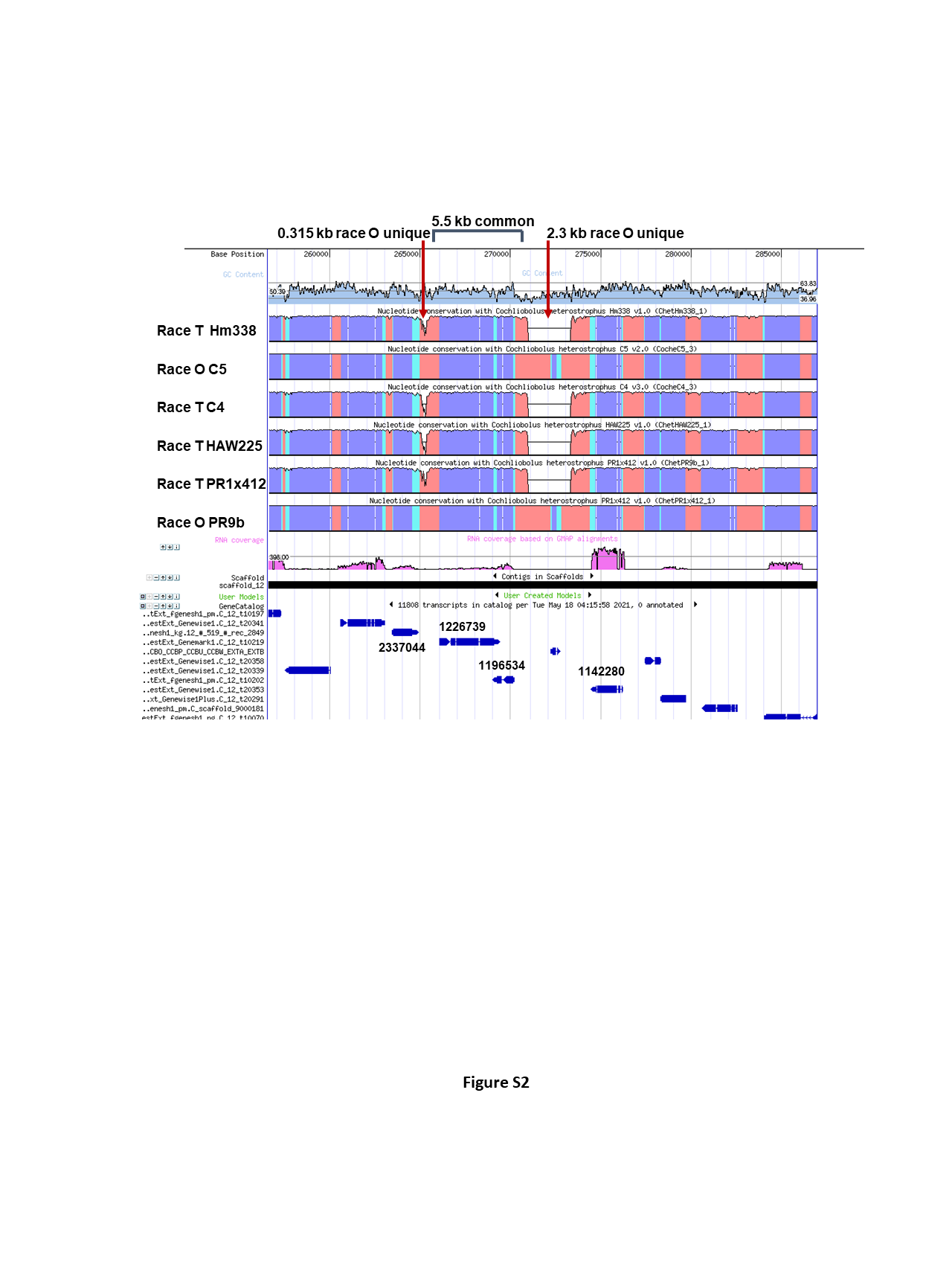

### Suppl Fig 3

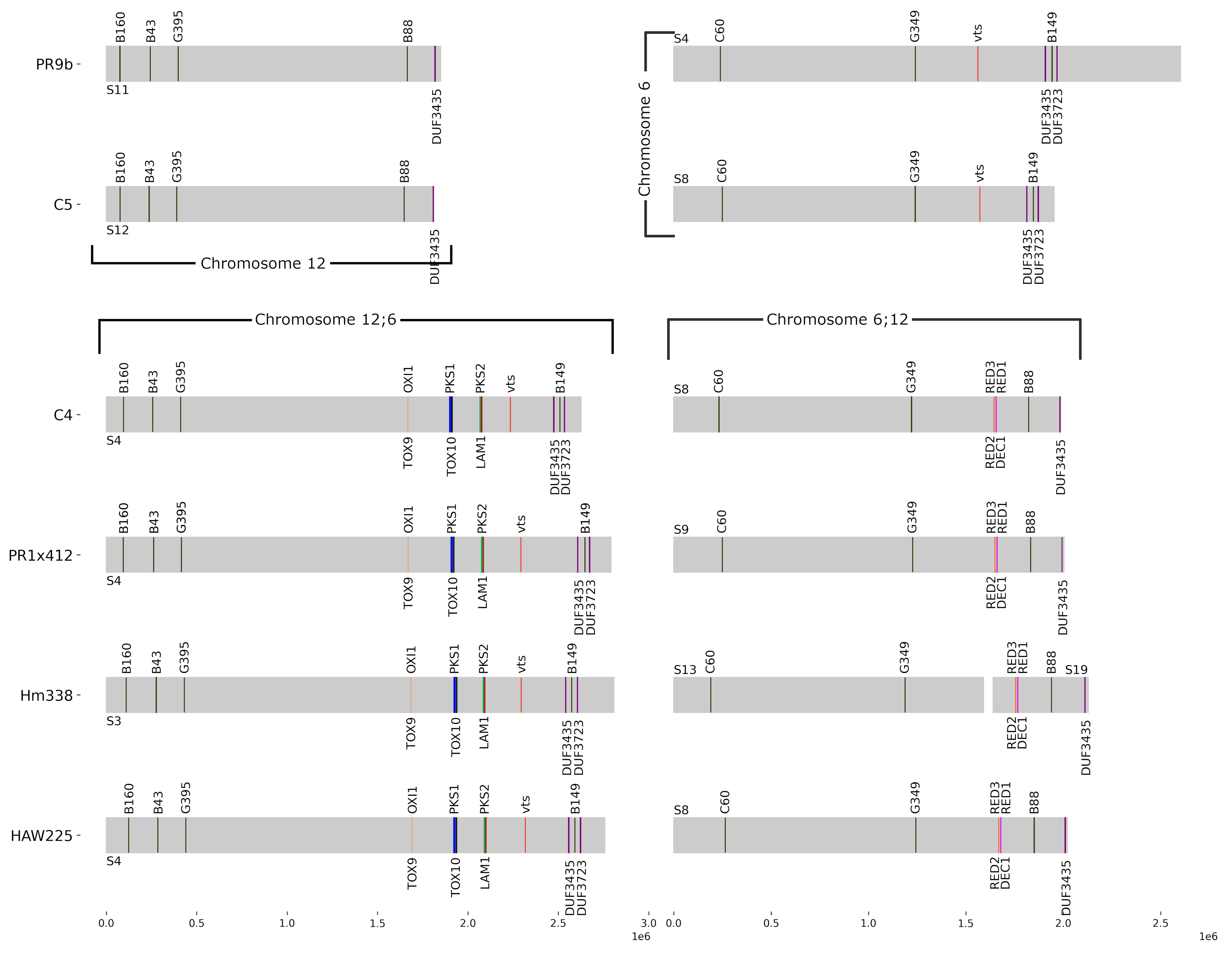

### Suppl Fig. 1

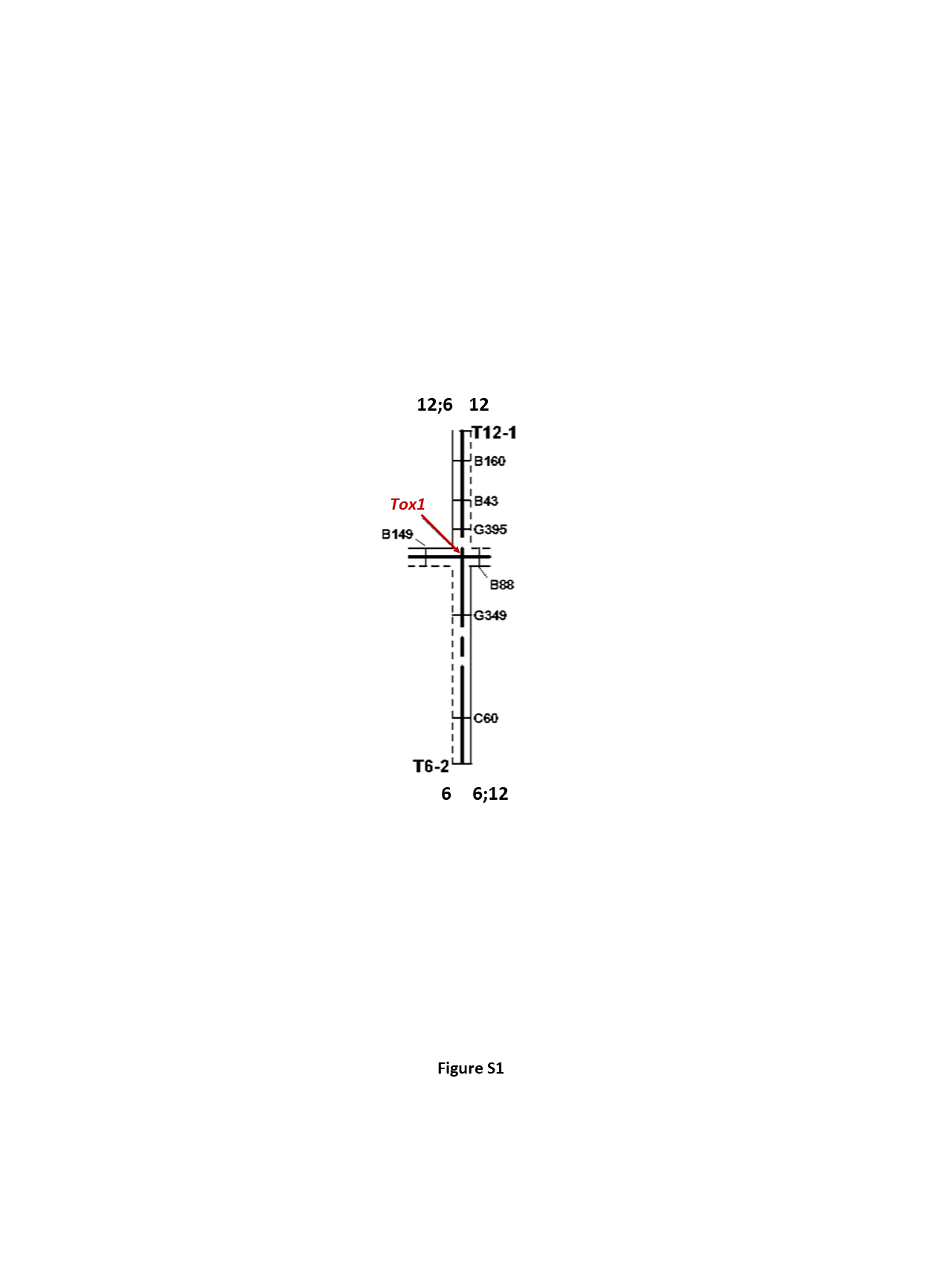
